# Effects of different load on physiological, hematological, biochemical, cytokines indices of Zanskar ponies at high altitude

**DOI:** 10.1101/262253

**Authors:** Prince Vivek, Vijay Kumar Bharti, Deepak Kumar, Rohit Kumar, Kapil Nehra, Dhananjay Singh, Om Prakash Chaurasia, Bhuvnesh Kumar

## Abstract

High altitude people required high endurance pack animals for load carrying and riding at prevalent mountainous terrains and rugged region. So far no studies have been taken to evaluate effect of loads on physiology of ponies in high altitude region. So, in this view we evaluated variation in physiological, hematological, biochemical, and cytokines indices of Zanskar ponies during load carrying at high altitude. Total twelve (12) numbers of Zanskar ponies, mare, age 4–6 years, were divided into three groups; group-A (without load), group-B (60 kg), and group-C (80 kg) of back pack loads. Track was very narrow and slippery with gravel, uneven with rocky surface and has a steep gradient of 4 km uphill at altitude 3291 to 3500 m. When we evaluate these parameters, it is understood that the heart rate, pulse rate and respiration rate was significantly *(p<0.05)* increased in 80 kg group among the three groups. The hematology parameters viz. hemoglobin, PCV, lymphocytes, monocytes%, ESR and eosinophil% significantly *(p<0.05)* changed in 80 kg group after load carrying among the three groups which was followed by control and 60 kg group. In biochemical parameters viz. LA, LDH, TP, HK, CORT, T_3_, CRT, AST, CK-MB, GPx, FRAP and IL-6 significantly *(p<0.05)* changed in 80 kg group after load carrying among the three groups which was followed by control and 60 kg group. The ALT, ALB, GLB, UR and UA significantly *(p<0.05)* changed in 80 kg group before and after load carrying among the three groups which was followed by control and 60 kg group. It has been concluded that, this result has revealed strong correlation of change in biomarkers level with performance in ponies during load carry. Hence, these parameters might be use for performance of endurance of Zanskar ponies in high mountain region.

## INTRODUCTION

The strategic importance of protecting high mountain regions to prevent hostile infiltration requires huge deployment of troops along with logistic supply chain. Equine have been playing significant role in logistic support (Bharti et al., 2017). Even in today’s era of modernization the equine draught power cannot be wished away for many strategic reasons. L-Sector (Ladakh sector), prevalent high altitude region is a very difficult rugged terrain with highest peaks in the world, where large number of people working for recreation. The prevalent mountainous terrains and rugged region are not suitable for motorized vehicle to use in logistic transport. Therefore, people required high endurance pack animals for load carrying and riding. Therefore, there is a need of huge requirement of logistic support to deploy as an all terrain vehicles in mountainous areas. These animals have been virtually indispensable to human civilisation and are widely use for load carrying and transportation purposes by local people of remote localities and rough terrains of high mountains. The whole mechanics of ponies is much suitable and ideal for balancing while riding and load carrying. Hence, due to their suitability as pack animals, ponies are in high demand for deployment as all terrain vehicles in L-sector and other high mountain part of country (Venkatesan et al., 2011).

The change in physiological, hematological and biochemical blood analysis provides information of clinical health status of the animals (Burlikowska et al., 2015). Different types of activities viz. exercise, intensity, duration and frequency influencing these parameters in horses (Ciesla et al., 2013). These indexes assume much more significance in horses owing to the use as the indicator of their physical fitness and performance (Snow, 1985). Stress acting on body can disturbed the homeostatic balance (Cuesta and Singer’s, 2012). Fitness or exercise tolerance of a horse can be determined by assessment, through physical examination, of heart and respiratory rates (Bashir and Rasedee, 2009). Endurance exercise is one of the best exercise programs to asses a safe and effective fitness levels in athletes as well as in pack animals carrying endurance load. A healthy cardiac system is must for high level of fitness and wellness of working pony. The cardiovascular system can maintain hemodynamic homeostasis in response to repetitive environmental stressor by increasing its functional reserve capacity (Vivek et al., 2018). So, cardiac endurance performance is a key of health assessment of fitness level during performing endurance exercise and load carrying. An example of repetitive physiologic stress is endurance exercise training which induces adaptation in the cardiovascular system (heart) characterized by increase in maximal cardiac output, stroke volume, diastolic filling and left ventricular volume overload hypertrophy (Scharhag et al., 2002). These adaptative responses are in concert with the adaptation in skeletal muscles (Holloszy and Coyle, 1984) which increase maximal oxygen (O2) uptake during training. In veterinary medicine and physiology, there has been a need to discover novel cardiovascular biomarkers to aid in the early detection, diagnosis, and prognosis of high altitude disease in animals (Bharti et al., 2016). However, to increase their load carrying performance we need regular up-gradation and culling of weak animals having poor genetic make-up in endurance performance. Therefore, there is need a need to identify biomarkers of endurance performance and load carrying for selection of high performing breedable pack animals and deployment of high load carrying animals to reduce the maintenance cost and loss of animals on deployment which further necessitates the need of research and development to identify biomarkers at high altitude in ponies.

**Fig. 1.**
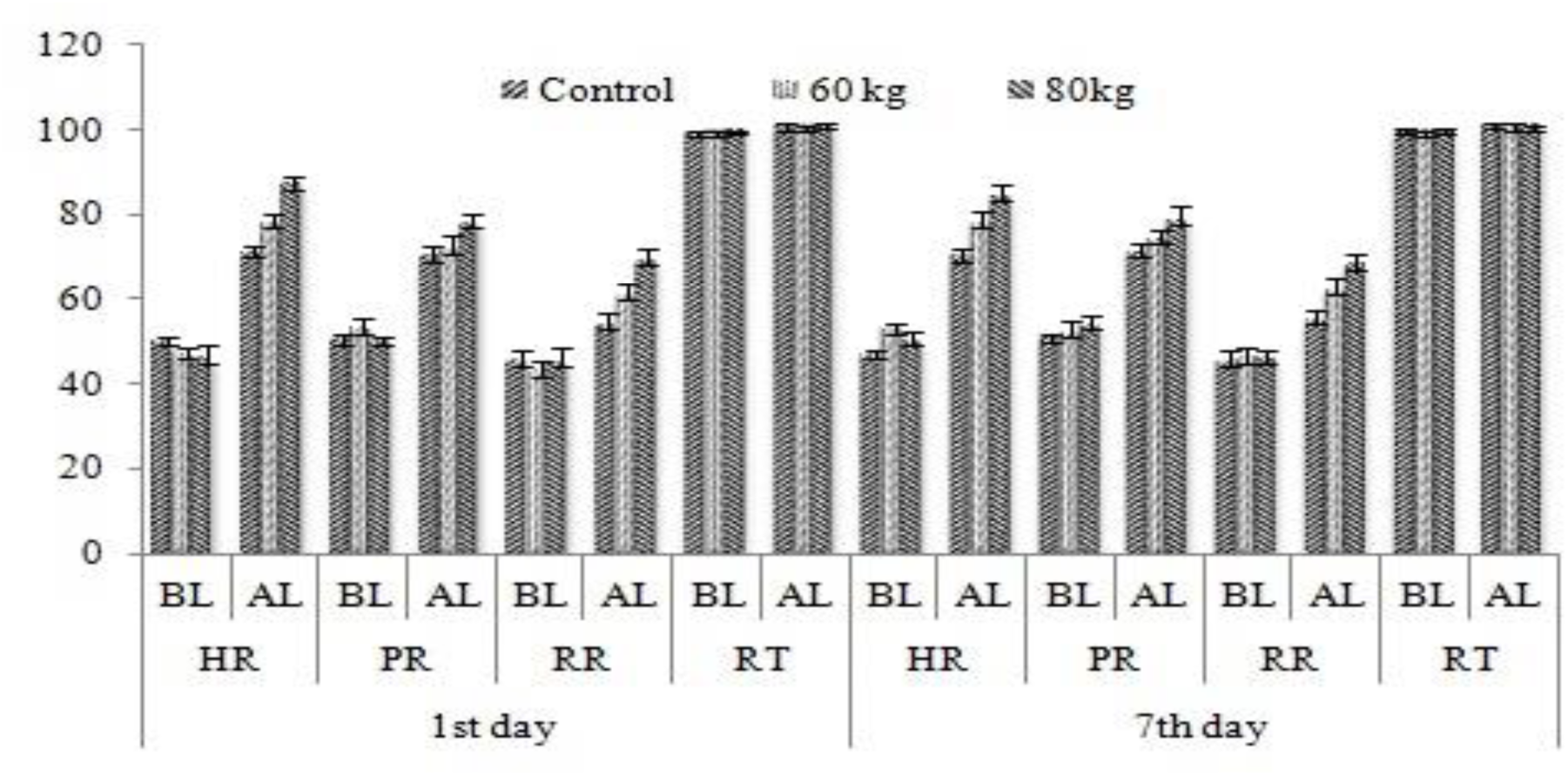
Change in physiological responses during load carry of Zanskar pony at high altitude *Values are presented as mean ± SEM; *indicate significant difference (p<0.05) within the graph in comparison with Group-A with Group B & C.* BL-Before load, AL-After load; Physiological responses *viz*. HR**-** Heart rate (beat/min); PR-Pulse rate (pulse/min), RR-Respiration rate (cycle/min); RT-Rectal temperature (°F).

**Fig. 2–5.**
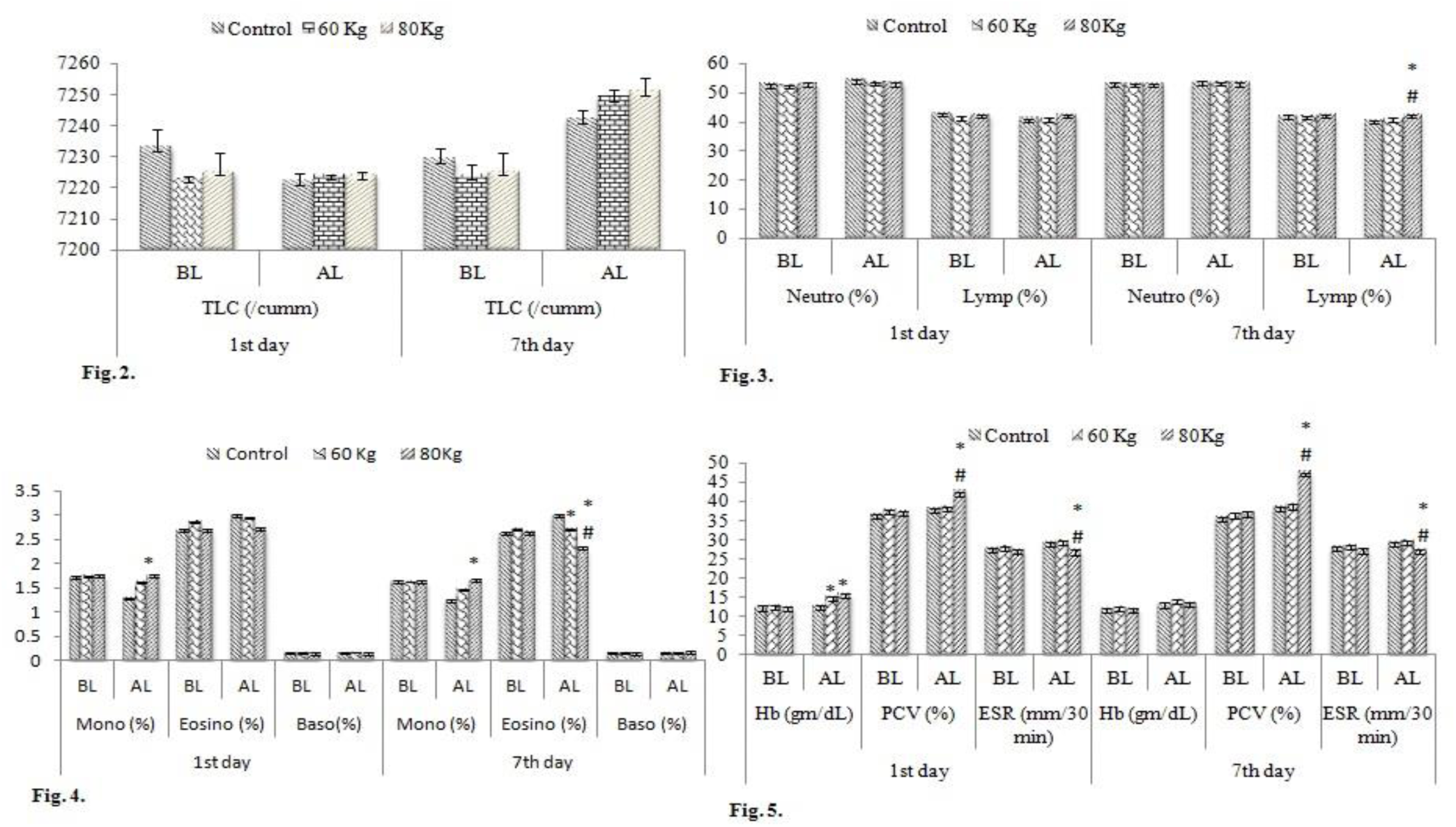
Change in hematological parameters during load carry of Zanskar pony at high altitude *Values are presented as mean ± SEM; *# indicate significant difference (p<0.05) within the graph in comparison with Group-A with Group B & C.* BL-Before load, AL-After load; Hb-Hemoglobin; PCV-Packed cell volume; ESR-Erythrocyte sedimentation rate; TLC-Total leckocyte count; Neutro-Neutrophil; Lymp-Lymphocyte; Mono-Monocyte; Eosino-Eosinophil; Baso-Basophil

**Fig. 6–13.**
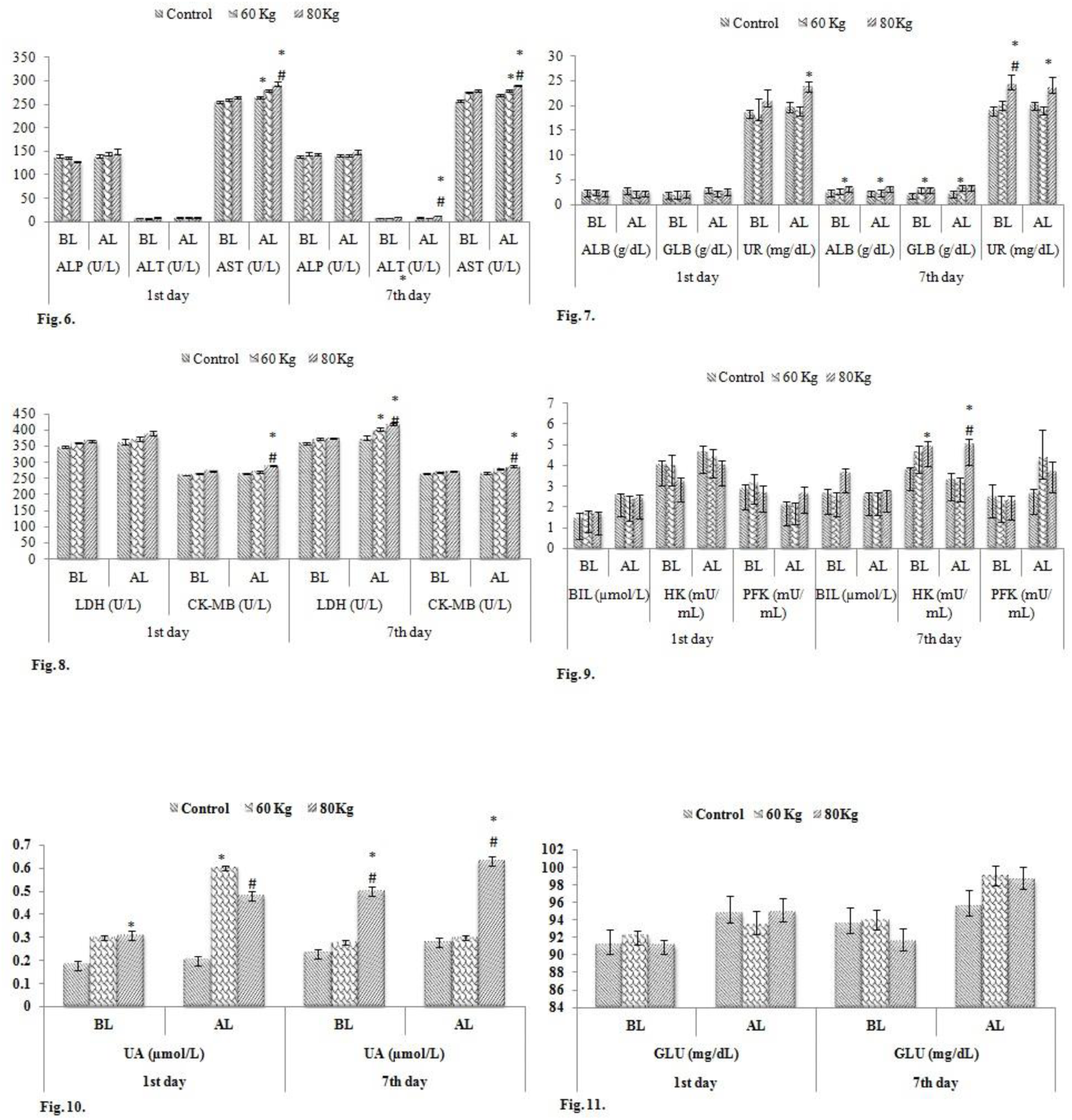
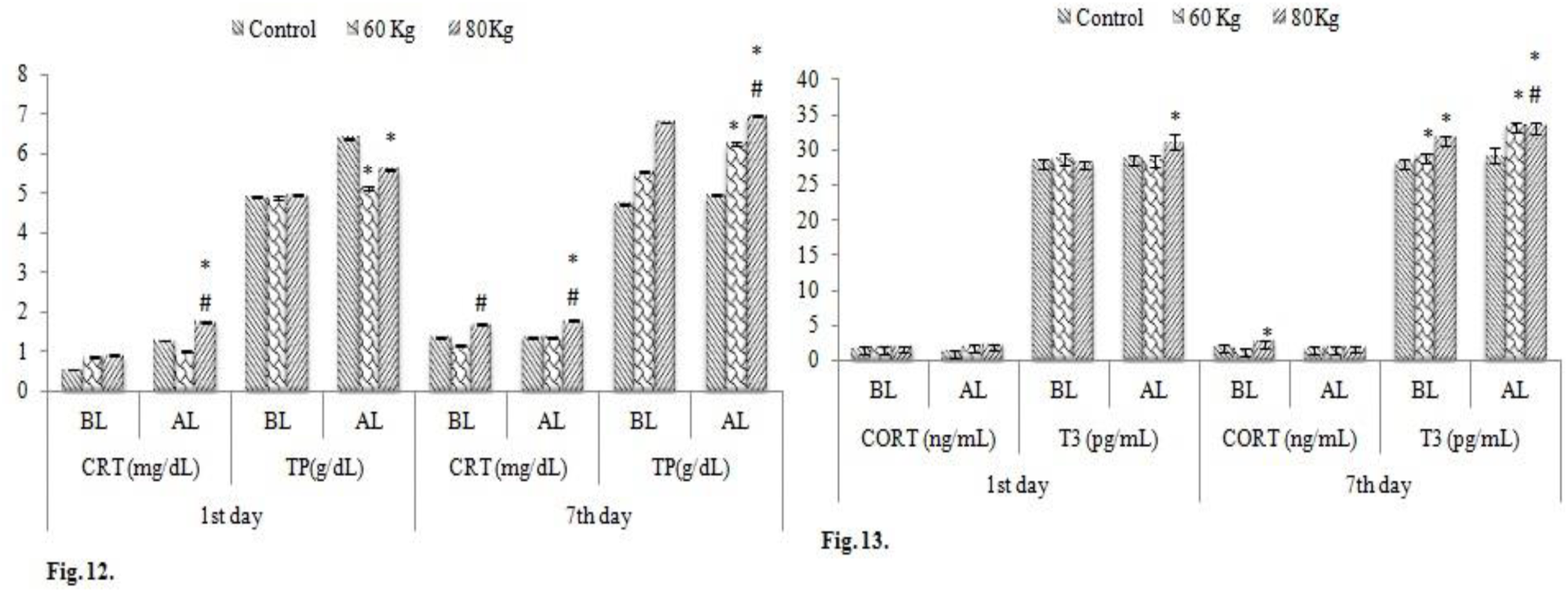
Change in patho-biochemical biomarkers during load carry of Zanskar pony at high altitude *Values are presented as mean ± SEM; *# indicate significant difference (p<0.05) within the graph in comparison with Group-A with Group B & C.* BL-Before load, AL-After load; ALP: Alkaline phosphatase; ALT: Alanine aminotransferase; AST: Aspartate aminotransferase; LDH: Lactate dehydrogenase; CK-MB: creatine kinase isotype; ALB: Albumin; GLB: Globulin; UR: Urea; BIL: Bilirubin; HK: Hexokinase; PFK: Phosphofructokinase; CORT: Cortisol; T_3_: Triiodothyronine; GLU: Glucose; CRT: Creatinine; TP: Total protein; UA: Uric acid.

**Fig. 14–15.**
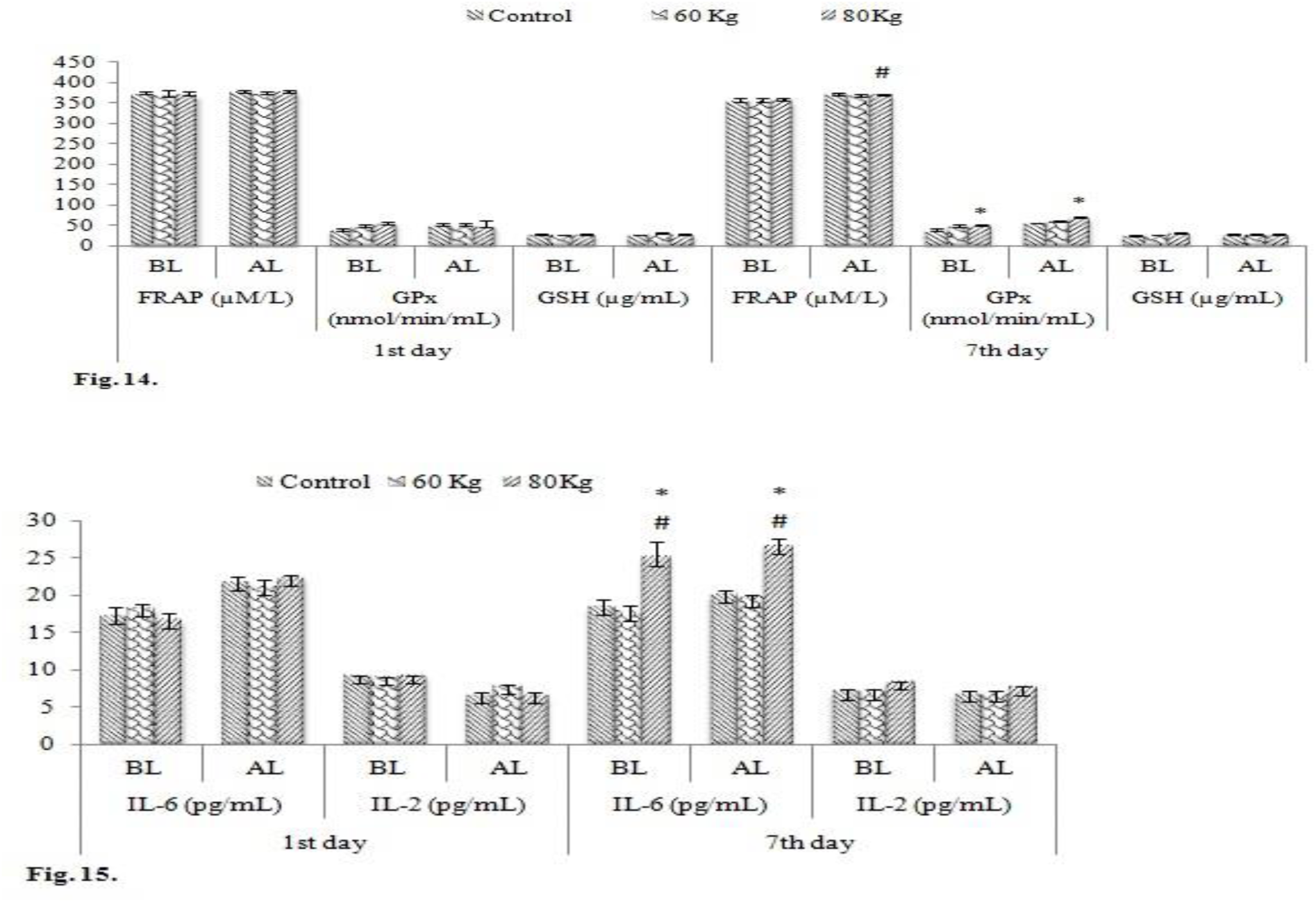
Change in patho-biochemical, oxidative stress, antioxidant and cytokine biomarkers during load carry of Zanskar pony at high altitude *Values are presented as mean ± SEM; *# indicate significant difference (p<0.05) within the graph in comparison with Group-A with Group B & C.* BL-Before load, AL-After load; FRAP: Ferric reducing antioxidant power; GPx: Glutathione peroxidase; GSH: Total glutathione; IL-6: Interleukin-6; IL-2: Interleukin-2

## RESULTS

The heart rate, pulse rate and respiration rate increased significantly *(p<0.05)* in group C (80 kg) which was followed by group-A (control) and group-B (60 kg) for throughout the experiment after load carrying. The hematology parameters viz. hemoglobin increased significantly *(p<0.05)* in 80 kg group after load carrying on 1^st^ day among the three groups which was followed by control and 60 kg whereas, PCV, lymphocytes, monocytes percentage also increased significantly *(p<0.05)*. However, erythrocyte sedimentation rate (ESR) and eosinophil percentage decreased significantly *(p<0.05)* in 80 kg group after load carrying on 7^th^ day among the three groups which was followed by control and 60 kg group. In biochemical parameters viz. LA, LDH, TP, HK, CORT, T3, GPx, FRAP and IL-6 significantly *(p<0.05)* increased in 80 kg group on 7th day after load carrying among the three groups which was followed by control and 60 kg group. The ALT, ALB, GLB, UR and UA increased significantly*(p<0.05)* in 80 kg group on 7^th^ day before and after load carrying among the three groups which was followed by control and 60 kg group. The CRT, AST and CK-MB were increased significantly *(p<0.05)* in 80 kg group on 1st and 7th day after load carrying among the three groups which was followed by control and 60 kg group.

## DISCUSSION

The physiological response viz. heart rate, pulse, respiration rate and rectal temperature increased significantly after load carry in animals carrying more weight comparatively; these changes were associated with either the level of load or the changes agreement as per reported finding by (Stefansdottir et al., 2017) in horses carrying a rider of different weight. It is well known that respiration rate is somewhat limited by stride frequency during exercise and also depend on gait pattern (Butler et al., 1993) therefore breathing can do more of a catch-up post exercise. As per our knowledge little is known about how respiration rate is affected by comparatively carrying more loads at high altitude. It may assist in cooling the body during carrying load, although vasodilation and sweating are more important (Hodgson, 2014). Nonetheless, the rectal temperature of the ponies was increased indicating a substantial need for cooling.

Hemoglobin and PCV concentration increased after carrying more weight comparatively in the ponies as the amount of red blood cells released from the spleen reflects the magnitude of load until maximum splenic contraction. While packed cell volume can predict the oxygen transportation capacity. These changes are in accord with the earlier finding reported by (Stefansdottir et al., 2014) in horses, their riding assessment and by (Kang and Park, 2017) in horses after endurance exercise. These parameters grant us meaningful information about fitness changes in horses (Piccione et al., 2010). The erythrocyte sedimentation rate (ESR) was decreased before and after in comparatively more load carrying ponies, which as the reported finding reported by (Hiraga and Sugano, 2016) during the intense exercise. It was observed that more weight load carrying ponies showed increase in lymphocytes percentage this evoke in leukocytosis, stayed in agreement with the results of other studies (Janick et al., 2013). The reaction of exercise-induced release of lymphocytes to peripheral blood was caused by increased levels of catecholamines, especially epinephrine (Iversen et al., 1994). Monocytes were found to be increased after load carrying, which was in agreement with the results of study (Robson et al., 2003). The strongest point of this study is the fact that it was the first to study to analyze the effects of different load carrying at high altitude on hematological indices viz. lymphocytes, monocytes and eosinophil, hemoglobin and PCV in native breed of ponies.

Interestingly, we found that biochemical metabolites and their enzymes viz. lactic acid increased after load carrying in the high pack group, these results indicates that ponies with more muscular work either had more support from aerobic muscle performance or that their muscle tissue was larger or better at metabolizing lactate during load carrying. Plasma lactate levels exceed 4 mmol/l can only be maintained for relatively short periods (minutes) without causing fatigue or reducing performance (Stefansdottir et al., 2017). It was suggested that the higher response of glycolytic metabolism might be reflected in higher values of CK and AST because of changes in permeability of the muscle fibre membranes (Fazio et al., 2011). The increase in ALT level after carrying more weight in these ponies addressed the question regarding the potential changes in ALT level post-exercise. It is reasonable to assume, that measurements of ALT levels might be reflected by load carrying programs in ponies at high altitude and impact on muscle mass because there is a strong correlation between AST and ALT, as well as between CK and ALT (Greco-Otto et al., 2017). Recently one of the study observed that low ALT blood activity, as a biomarker for increased weakness, is associated with lower baseline fitness of post-acute myocardial infarction and post-cardiac surgery patients going through a cardiac rehabilitation program. Low level of ALT serves as an independent predictor for low baseline exercise capacity, it is also independent from other physiologic variables that share this prognostic value by themselves including blood haemoglobin concentration and left-ventricular heart failure parameters (Kogan et al., 2018). In light of our findings, it might be logical to suggest that ALT may be used, to identify endurance biomarkers in ponies at high altitude (Vivek et al., 2018). Several studies confirm that at high alanine concentrations, ALT-mediated is the main step which controls the gluconeogenic flux from alanine (Groen et al., 1982, Burelle et al., 2000). ALT is not a specific marker to liver cells, but also a marker of damages in other tissues, including skeletal muscle, which releases these enzymes and leads to elevated levels (Vrankovic et al., 2015). A significant increase in AST value after the jumping test suggests increase in mitochondrial membrane permeability, rather than muscle damage or strenuous muscular exercise (Valberg, 2009). The LDH activities are considered to be the main indicators of myocyte stability. The elevated level of lactate dehydrogenase is the result of more anaerobic energy supply in the body (Vrankovic et al., 2015). Studies of enzymatic patterns are helpful in determining the degree of muscles, heart and liver adaptation to endurance exercise and load carrying. The level of total protein, albumin, globulin, creatinine and urea were found to be significantly increased. Greco-Otto et al., 2017 noticed that there was a strong correlation between plasma total protein, albumin, globulin, creatinine and urea. This increase in the values of these parameters was due to dehydration during intense work (Munoz et al., 2011). An increased concentration of urea has been associated with liver function and has been suggested to indicate increased synthesis of urea from ammonia in liver. The association of increased level of urea and creatinine with a well prognosis may suggest controlled diuresis and probably to inhibit liver failure (David Weiner, 2014). Increased concentration of uric acid acts as a potent scavenger of ROS and acts like antioxidant and protects from oxidative stress during intense work. Several published investigations of high altitude report a rise in circulating plasma uric acid levels (Finaud et al., 2006; Sinha et al., 2009; Peters et al., 2015). Creatinine is produced by the decomposition of creatine, a nitrogen compound used by muscle cells to store energy. The serum concentration of creatinine varies according to creatine synthesis and the amount of muscle tissue of that animal (Stockham, 1995). Increased HK activity leads to an enhanced capacity for phosphorylating exogenous glucose (Susana et al., 2006). CK-MB enzyme activity increased after load carry because of structural damage of muscles during load carrying. Our data agrees well with the previous investigations on ultra-marathon run (Son et al., 2015). Past studies have suggested that the increased level of CK-MB helps transfer of high energy phosphates into and out of mitochondria, thus helping in regeneration of ATP in the cells (Mannem et al., 2009). The T_3_ increased after load carry because it was primarily related to their involvement in maintaining functional homeostasis and hence increases metabolic activity, which helps burn more calories. According to previous studies T_3_ was reported to increase in regular exercise group (Bansal et al., 2015). The concentration of cortisol increased after more weight load carry in ponies because of increased oxidative stress which can initiate a host of metabolic changes such as increasing gluconeogenesis and decreasing the amount of glucose that is metabolised as it is considered as stress as well as a catabolic hormone. Cortisol can also stimulate lipolysis in adipocytes which increases the release of lipids (Viru and Viru, 2001; Kramer and Ratamess, 2005). The FRAP value increases after comparatively more weight load carrying ponies which is in accord with the study done before by (Balogh et al., 2001), and stated that the rise in FRAP values in these horses was probably caused by increased uric acid concentration. The increased level of GPx activity after weight load carrying ponies are in agreement with the study of (Wieslaw, 2010), and the demonstrated that among the antioxidants, the activity of GPx showed the high post-exercise changeability, which suggests a great importance of this enzyme in the protection of the organism from the increased generation of reactive oxygen species and suggested that the values of this index most strongly correlate with GPx activity, which may suggest its usefulness in determining the antioxidative potential in horses. Increase in circulating levels of IL-6 after load carry are in agreement with the study reported by (Steensberg et al., 2000), and they stated that IL-6 increases after exercise even without muscle damage. This result suggests that increased IL-6 levels in higher load carrying ponies will use more fat as energy than before load carrying resulting in increased lypolysis. Moreover, past evidence stated that in responses to exercise IL-6 acts like a myokine which is synthesized in skeletal muscle and is secreted into the bloodstream and during muscular activity has a great potential to stimulate lipolysis and provides energy to working muscle (Grazyna Lutoslawska, 2012). Plasma IL-6 increases in an exponential fashion with exercise and is related to exercise intensity, duration, the mass of muscle recruited, and their endurance capacity (Pedersen and Hoffman, 2000). The interesting point of this study is to assess the circulating levels of cytokines in different load carrying ponies at high altitude.

## CONCLUSION

It has been concluded that, heart rate, respiration rate, hematological indices like PCV, lymphocytes, monocytes, Hb and ESR, biochemical indices like lactic acid, LDH, TP, HK, CORT, T3, ALT, AST and CRT, ALB, GLB, UR, UA, GPx, FRAP and IL-6 are important biomarkers to asses effect of load on animal physiology and endurance. Further, this result has revealed strong correlation of change in biomarkers level with performance in ponies during load carry. Hence, these parameters might be use for performance of endurance of Zanskar ponies in high mountain region.

## MATERIALS AND METHODS

This experiment was conducted at Zanskar ponies breeding unit located at Army RVS unit in Pratapur (Nubra vally) for 7 day on a track which had a steep climb of 4 km uphill, the uneven rocky surface, at an altitude of 3291 m to 3500 m (end point) was made of gravel and it was slippery. The ambient temperature was in the range of 15–30 ⁰C. These animals were divided into three groups n=04 in each group, Group-A as control (without load), Group-B (60 kg) and Group-C (80 kg) carrying the specified back pack loads including the weight of saddle. Ponies were made to walk on hill before sampling to ensure their wellness and health. Their gait was closely observed to rule out ponies exhibiting any difficulties during performance of the task. This practice helped us in ensuring that all the selected ponies are well coordinated and are equally able to perform the task comfortably. Their load carrying ability was standardize before the experimental trial.

### Parameters studied

The physiologic responses viz. heart rate (HR), pulse rate (PR), respiration rate (RR), and rectal temperature (RT) were recorded before and immediately after the load carry. A total of 15 ml blood sample were collected from the jugular vein in vacutainer tubes just before and immediately after the load carry on 1^st^ and 7^th^ day for analysis of hematological parameters. Total leukocyte counts (TLC) were determined by haemocytometer methods. Hemoglobin (Hb) concentration was determined by using Sahli’s apparatus, and packed cell volume (PCV) by microhaematocrit. Erythrocyte sedimentation rate (ESR) was determined by using Wintrobe Haematocrit graduated tubes (Benjamin 1978). Differential leukocyte count (DLC) was obtained by the method of Kolmer et al. (1961).

However, the serum isolated on 1^st^ and 7^th^ day was used for analysis of biochemical, oxidative stress and cytokine indices. The biochemical parameters were viz. alkaline phosphatase (ALP), alanine aminotransferase (ALT), aspartate aminotransferase (AST); lactate dehydrogenase (LDH), creatine kinase isotype (CK-MB), glucose, total protein (TP), albumin (ALB), urea (UR) and uric acid (UA) were determined by using commercial kits and by using fully automated BS-120 clinical biochemistry analyzer as per the manufacturer’s protocol given in product brochure. Serum triiodothyronine (T_3_) and cortisol (CORT) were estimated by using commercial ELISA kits as per the manufacturer’s protocol. Hexokinase (HK) and Phosphofructokinase (PFK) were estimated by using colorimetric assay kit as per the manufacturer’s protocol. whereas, lactic acid (LA) assay was performed as per Pryce method (1969) by the help of molecular device SpectraMax^R^ i3x.

The oxidative stress parameters in serum samples were measured by Ferric reducing antioxidant power (FRAP) assay as demonstrated by Benzie and Strain, 1996 with slight modifications. Serum glutathione peroxidase (GPx) level was measured through colorimetric method whereas reduced glutathione (GSH) levels were measured through the fluorometric method as per the manufacturer protocol by the help of molecular device SpectraMax^R^ i3x. The cytokine indices viz. IL-2 and IL-6 were estimated by using commercial ELISA kits as per the manufacturer’s protocol with the help of molecular device SpectraMax^R^ i3x.

### Statistical analysis

With the help of Graph Pad Prism5, data were analysed by one-way ANOVA method and level of significance considered at *p* < 0.05. Results are expressed as mean ± SEM (standard error mean).

## Acknowledgments

This study was supported in the form of a Senior Research Fellowship (SRF) to the first author (P. Vivek) by CSIR-HRDG, New Delhi and research facilities provided by Defence Institute of High Altitude Research (DIHAR), DRDO, India for conducting this study is duly acknowledged. We would like to thank all the technical staff of our animal facility and Army’s personnel (RVC) for the husbandry care of the ponies and assistance during sampling.

## Authors’ Contribution

Authors contributed to study design, data collection, data analysis, interpretation and manuscript preparation.

## References

Bharti, V. K., Vivek, P., Arora, A., Balaje, S. S. and Chaurasia. O. P. (2017). Cardiovascular biomarkers of endurance performance in ponies: Selection aid for high load carrying pack animals under high altitude stress condition. Society of Animal Physiologists of India, XXVI Annual Conference.

Burlikowska, K., Boguslawska-Tryk, M., Szymeczko, R. and Piotrowska, A. (2015). Haematological and biochemical blood parameters in horses used for sport and recreation. JCEA 16(4).

Benjamin, M. M. (1978). Outline of veterinary clinical pathology. Iowa State University Press.

Bashir, A. and Rasedee, A. (2009). Plasma catecholamines, sweat electrolytes and physiological responses of exercised normal, partial anhidrotic and anhidrotic horses. Am. J. Anim. Vet. Sci. 4, 26–31.

Benzie, I. F. and Strain, J. J. (1996). The ferric reducing ability of plasma (FRAP) as a measure of “antioxidant power”: the FRAP assay. Anal. Biochem. 239, 70–76.

Butler, P. J., Woakes, A. J., Smale, K., Roberts, C. A., Hillidge, C. J., Snow, D. H. and Marlin, D. J. (1993). Respiratory and cardiovascular adjustments during exercise of increasing intensity and during recovery in thoroughbred racehorses. J. Exp. Biol. 179, 159–180.

Burelle, Y., Fillipi, C., Péronnet, F. and Leverve, X. (2000). Mechanisms of increased gluconeogenesis from alanine in rat isolated hepatocytes after endurance training. Am. J. Physiol. Endocrinol. Metab. 278, E35–42.

Bansal, A., Kaushik, A., Singh, C. M., Sharma, V. and Singh, H. (2015). The effect of regular physical exercise on the thyroid function of treated hypothyroid patients: An interventional study at a tertiary care center in Bastar region of India. AMHS. 3, 244.

Balogh, N., Gaal, T., Ribiczeyne, P. S. and Petri, A. (2001). Biochemical and antioxidant changes in plasma and erythrocytes of pentathlon horses before and after exercise. Vet. Clin. Pathol. 30, 214–218.

Ciesla, A., Pilarczyk, B., Tomza-Marciniak, A., Pikula, R. and Smugala, M. (2013). Effect of the intensity of recreational horse use in the summer holiday season on serum Se concentration and chosen haematological parameters. Acta. Sci. Pol. Zootechnica. 12(3).

Cuesta, J. M. and Singer, M. (2012). The stress response and critical illness: a review. Crit. Care. Med. 40, 3283–3289.

Coffey, V. G. and Hawley, J. A. (2007). The molecular bases of training adaptation. Sports med. 37, 737–763.

de Paula Nogueira, G., Barnabe, R. C., Bedran-de-Castro, J. C., Moreira, A. F., Fernandes, W. R., Mirandola, R. M. and Howard, D. L. (2002). Cortisol serico, concentraçao de lactato e creatinina em cavalos de corrida Puro Sangue Inglês com diferentes idades e estágios de treinamento. Braz. J. Vet. Res. An. Sci. 39, 54–57.

Fazio, F., Assenza, A., Tosto, F., Casella, S., Piccione, G. and Caola, G. (2011). Training and haematochemical profile in Thoroughbreds and Standardbreds: A longitudinal study. Livest. Sci. 141, 221–226.

Finaud, J., Lac, G. and Filaire, E. (2006). Oxidative stress. Sport. Med. 36, 327–358.

Groen, A. K., Sips, H. I., Vervoorn, R. C. and Tager, J. M. (1982). Intracellular compartment ation and control of alanine metabolism in rat liver parenchymal cells. FEBS J. 122, 87–93.

Giri, A., Bharti, V. K., Kalia, S., Kumar, B. (2016). Cardiovascular biomarkers of high altitude adaptation: Selection aid for livestock breeding. Int. J. Bioassays. 5, 5146–5150.

Greco-Otto, P., Massie, S., Shields, E., Roy, M. F., Pajor, E. and Leguillette, R. (2017). High intensity, short duration pulling in heavy horses: physiological effects of competition and rapid weight change. BMC Vet. Res. 13, 317.

Hodgson, D. R., McKeever, K. H. and McGowan, C. M. (2014). The athletic horse: principles and practice of equine sports medicine. Elsev. Health. Sci.

Hiraga, A. and Sugano, S. (2016). Studies on exercise physiology and performance testing of racehorses performed in Japan during the 1930s using recovery rate as an index. J. Equine. Sci. 27, 131–142.

Iversen, P. O., Stokland, A., Rolstad, B., Benestad, H. B. (1994). Adrenaline-induced leucocytosis: recruitment of blood cells from rat spleen, bone marrow and lymphatics. Eur. J. Appl. Physiol. Occup. Physiol. 68, 219–227.

Janicki, B., Kochowicz, A., Buzała, M. and Krumrych, W. (2013). Variability of selected clinical and haematological indices in young stallions during 100-day performance test. Bulletin of the Veterinary Institute in Pulawy. 57, 91–96.

Kolmer, J. A., Spulding, E. H. and Robinson, H. W. (1961) Approved laboratory technique, 5th edn. Published Place: 75.

Kang, O. D. and Park, Y. S. (2017). Effect of age on heart rate, blood lactate concentration, packed cell volume and hemoglobin to exercise in Jeju crossbreed horses. JAST. 59, 2.

Kogan, M., Klempfner, R., Lotan, D., Wasserstrum, Y., Goldenberg, I. and Segal, G. (2018). Low ALT blood levels are associated with lower baseline fitness amongst participants of a cardiac rehabilitation program. JESF. 16, 1–4.

Kraemer, W. J. and Ratamess, N. A. (2005). Hormonal responses and adaptations to resistance exercise and training. Sport. Med. 35, 339–361.

Krumrych, W.I. (2010). Blood antioxidant defence in horses during physical exercises. Bull. Vet. Inst. Pulawy. 54, 617–624.

Mannem, S. R., Ehtesham, M., Praveen, K. N. and Gupta, V. (2009). Elevated CK-MB without myocardial infarction. Sci. Med. 1(2).

Piccione, G., Messina, V., Casella, S., Giannetto, C. and Caola, G. (2010). Blood lactate levels during exercise in athletic horses. Comp. clin. Path. 19, 535–539.

Peters, B., Ballmann, C., Mcginnis, G., Epstein, E., Hyatt, H., Slivka, D., Cuddy, J., Hailes, W., Dumke C, Ruby B and Quindry J. (2016). Graded hypoxia and blood oxidative stress during exercise recovery. J. Sport. Sci. 34, 56–66.

Pedersen, B. K. and Hoffman-Goetz, L. (2000). Exercise and the immune system: regulation, integration, and adaptation. Physiol. reviews. 80, 1055–1081.

Robson, P. J., Alston, T. D. and Myburgh, K. H. (2003). Prolonged suppression of the innate immune system in the horse following an 80 km endurance race. Equi. Vet. J. 35, 133–137.

Shave, R., Howatson, G., Dickson, D. and Young, L. (2017) Exercise-Induced Cardiac Remodeling: Lessons from Humans, Horses, and Dogs. Vet. Sci. 4, 9.

Stefansdottir, G. J., Gunnarsson, V., Roepstorff, L., Ragnarsson, S. and Jansson, A. (2017). The effect of rider weight and additional weight in Icelandic horses in tolt: part I. Physiological responses. Animal. 1–9.

Sinha, S., Sing,h S. N. and Ray, U. S. (2009). Total antioxidant status at high altitude in lowlanders and native highlanders: role of uric acid. High alt. Med. & biol. 10, 269–74.

Sangiao-Alvarellos, S., Arjona, F. J., Míguez, J. M., del Río, M. P., Soengas, J. L. and Mancera, J. M. (2006). Growth hormone and prolactin actions on osmoregulation and energy metabolism of gilthead sea bream (Sparus auratus). Comp. Biochem. Physiol. Part A Mol. Integr. Physiol. 144, 491–500.

Steensberg, A., Hall, G., Osada, T., Sacchetti, M., Saltin, B. and Pedersen, B. K. (2000). Production of interleukin‐6 in contracting human skeletal muscles can account for the exercise‐induced increase in plasma interleukin‐6. J. Physiol. 529, 237–42.

Son, H. J., Lee, Y. H., Chae, J. H. and Kim, C. K. (2015). Creatine kinase isoenzyme activity during and after an ultra-distance (200 km) run. Biol. Sport. 32, 357.

Venkatesan, G., Deshmukh, P. B., Biswas, A., Bharti, V. K. and Srivastava, R. B. (2011). Zanskar Ponies: Packers and movers on the rugged terrain. In Innovations in Agro Animal Technologies, Editors: Srivastava R B, Selvamurthy W, Satish Serial Publishing House, Delhi, India. Pp. 234–245.

Vivek, P., Bharti, V. K., Giri, A., Kalia, S., Raj, T. and Kumar, B. (2018). Endurance exercise causes adverse changes in some hematological and physio-biochemical indices in Ponies under high altitude stress condition. Indian J. Anim. Sci. 88 (2), 91–00.

Vrankovic, L., Aladrovic, J., Beer-Ljubic, B., Zdelar-Tuk, M. and Stojevic, Z. (2015). Seasonal changes in enzyme activities and mineral concentrations in Holstein stallion blood plasma. Veterinarski arhiv. 85(3), 235–246.

Valberg, S. J. (2009). Heritable muscle diseases. Current Therapy in Equine Medicine 6461–6468.

Viru, A. A. and Viru, M. (2001). Biochemical monitoring of sport training. Human Kinetics.

Weiner, I. D., Mitch, W. E. and Sands, J. M. (2015). Urea and ammonia metabolism and the control of renal nitrogen excretion. Clin J Am Soc Nephrol. 10(8), 1444–1458.

